# Critical assessment of the ability of Boolean threshold models to describe gene regulatory network dynamics

**DOI:** 10.1101/2025.03.06.641948

**Authors:** Claus Kadelka, Kishore Hari

## Abstract

The inference of gene regulatory networks (GRNs) from high-throughput data constitutes a fundamental and challenging task in systems biology. Boolean networks are a popular modeling framework to understand the dynamic nature of GRNs. In the absence of reliable methods to infer the regulatory logic of Boolean GRN models, researchers frequently assume threshold logic as a default. Using the largest repository of published expert-curated Boolean GRN models as best proxy of reality, we systematically compare the ability of two popular threshold formalisms, the Ising and the 01 formalism, to truthfully recover biological functions and biological system dynamics. While Ising rules match fewer biological functions exactly than 01 rules, they yield a better average agreement. In general, more complex regulatory logic proves harder to be represented by either threshold formalism. Informed by these results and a meta-analysis of regulatory logic, we propose modified versions for both formalisms, which provide a better function-level and dynamic agreement with biological GRN models than the usual threshold formalisms. For small biological GRN models with low connectivity, corresponding threshold networks exhibit similar dynamics. However, they generally fail to recover the dynamics of large networks or highly-connected networks. In conclusion, this study provides new insights into an important question in computational systems biology: how truthfully do Boolean threshold networks capture the dynamics of GRNs?

**Significance statement:** Gene regulatory networks (GRNs) describe the complex interactions between genes. GRNs control biological processes, respond to environmental changes, and contribute to diseases. An accurate understanding of the dynamics of GRNs is therefore crucial. Given insufficient data, researchers frequently employ Boolean network models with default threshold logic to study the dynamics of GRNs. We systematically assess the ability of two commonly-used types of threshold networks to recover the true regulatory logic and the dynamics of published expert-curated Boolean GRN models. Inspired by biological insights, we further propose modifications to each threshold formalism that improve their match with biological networks.

## 1 Introduction

Gene regulatory network (GRN) inference, a fundamental but challenging task in computational systems biology, aims to uncover the complex interactions between genes from high-throughput expression data. Detailed knowledge of GRNs can help us understand how genes control biological processes, respond to environmental changes, or contribute to diseases. The complexity of GRN inference depends on the desired level of knowledge (Fig. 1). Earliest methods solely infer *undirected* co-expression networks [1,2]. Around 2010, the Dialogue on Reverse Engineering Assessment and Methods (DREAM) challenges boosted the development of numerous GRN inference methods, as well as their comprehensive assessment using both synthetic and real bulk transcriptomic data [3–5]. Many of these methods, e.g., the random forest-based GENIE3 [6], are able to infer *directed* networks. The advent of single-cell data led to the development of further, more specialized inference methods, comprehensively assessed in [7,8]. Some of these methods, e.g., SCODE [9] or SINCERITIES [10], tackle the even harder task of inferring *signed directed* networks, i.e., differentiating between positive (e.g., activation) and negative (e.g., inhibition) effects. Even a signed directed network is however static. Gene regulation, on the other hand, is a highly dynamic process.

**Figure 1.**
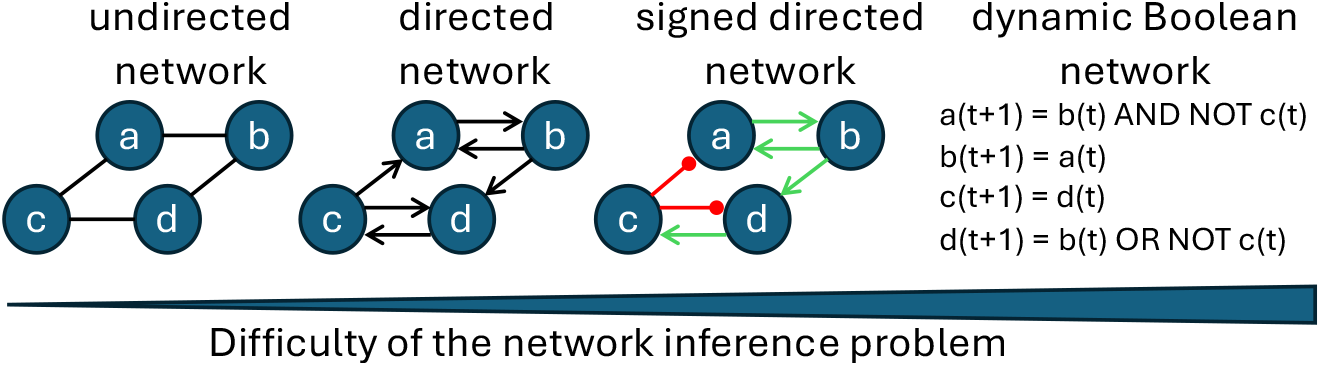
The difficulty of GRN inference depends on the desired level of knowledge. Given high-throughput transcriptomic data, the inference of undirected co-expression networks is the simplest task. Inferring the directionality of regulation and the type of regulation (positive: green arrows, negative: red arrows) adds complexity. Inference of the underlying Boolean regulatory logic is even much harder. This explains why frequently only the signed directed wiring diagram is known and default threshold assumptions are made about the regulatory logic.

**Figure 2.**
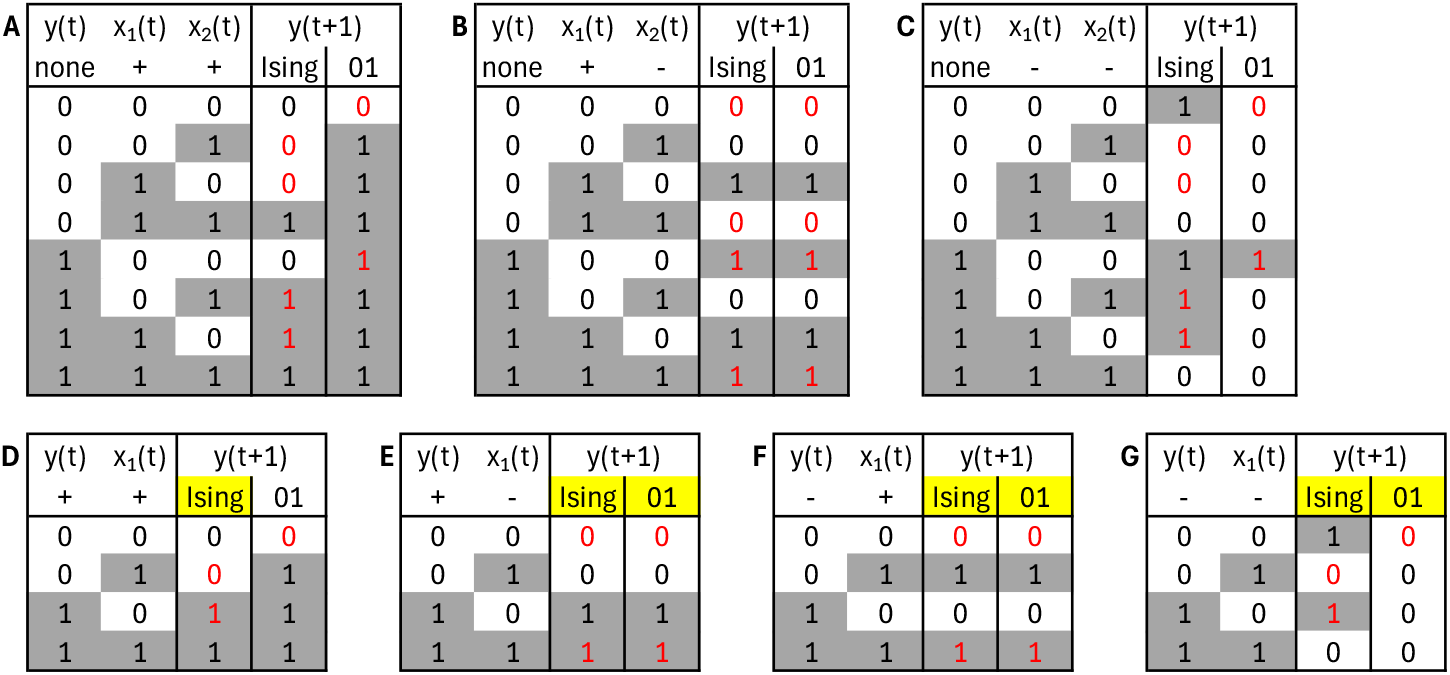
Boolean threshold functions with two inputs. Each table contains all combinations of Boolean inputs (left columns) and outputs under a different threshold formalism given a certain combination of activating (+) and inhibiting (−) inputs, as described in the second row. In (A-C), *y* does not regulate itself, i.e., the Boolean function is *y*(*t* + 1) = *f* (*x*_1_(*t*), *x*_2_(*t*)). In (D-G), *y* regulates itself, i.e., the Boolean function is *y*(*t* + 1) = *f* (*y*(*t*), *x*_1_(*t*)). Yellow cells highlight degenerated threshold functions (i.e., functions which do not depend on all inputs). Red entries indicate that the value of *y*(*t*) is used to determine *y*(*t* + 1).

**Figure 3.**
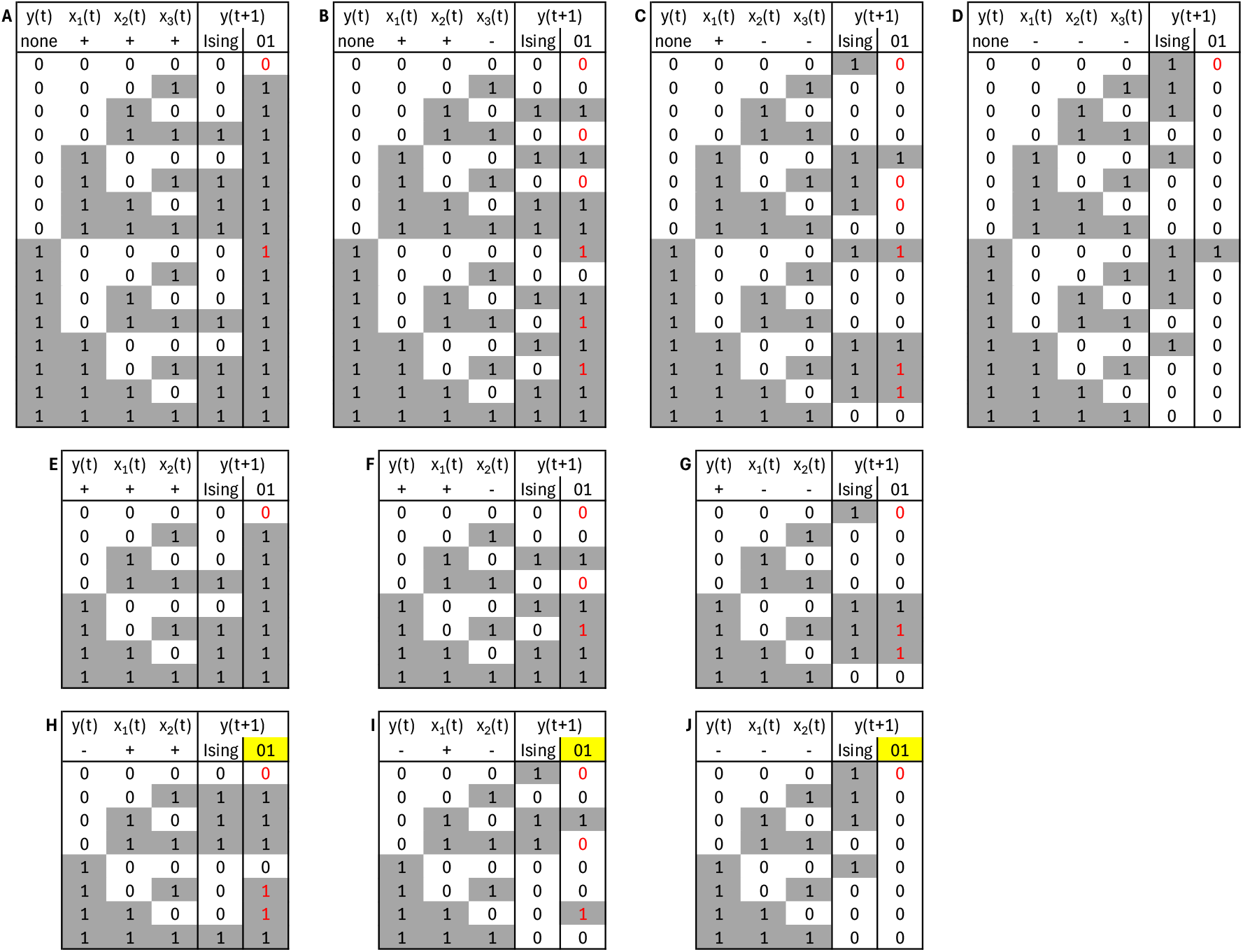
Boolean threshold functions with three inputs. Each table contains all combinations of Boolean inputs (left columns) and outputs under a different threshold formalism given a certain combination of activating (+) and inhibiting (−) inputs, as described in the second row. In (A-D), *y* does not regulate itself, i.e., the Boolean function is *y*(*t* + 1) = *f* (*x*_1_(*t*), *x*_2_(*t*), *x*_3_(*t*)). In (E-J), *y* regulates itself, i.e., the Boolean function is *y*(*t* + 1) = *f* (*y*(*t*), *x*_1_(*t*), *x*_2_(*t*)). Yellow cells highlight degenerated threshold functions (i.e., functions which do not depend on all inputs). Red entries indicate that the value of *y*(*t*) is used to determine *y*(*t* + 1).

Discrete dynamical systems (e.g., *Boolean networks*) are a popular tool to model the dynamic nature of GRNs. The static - frequently signed - directed graph, termed *wiring diagram* or *dependency graph*, describes the dependencies of the system. An additional set of update rules (e.g., Boolean functions) describes the regulatory logic governing the expression of each gene. In the simplest *Boolean* case, each gene is either on/active or off/inactive. Time is modeled in discrete time steps, and the expression level of a gene at a time step depends solely on the expression level of its regulators at the previous time step. Inference of the update rules represents a substantially more difficult task than inference of only the wiring diagram. Nevertheless, there exist several Boolean network inference tools (see e.g., [11,12] and a recent review [13]). To this day however, none of these tools can accurately and consistently infer complex Boolean GRNs.

To enable analyses of the dynamics of GRNs, modelers therefore rely on assumptions about the Boolean rules. A common assumption is that all rules are so-called *threshold rules*, where the number of active and inactive activators and inhibitors is compared to determine the expression of a gene at the next time point. By design, this assumes that gene regulatory processes are additive [14]. We know however that gene regulation often involves combinatorial interactions: A recent meta-analysis of 122 expert-curated Boolean GRNs found that the majority of gene regulatory logic is described by so-called *nested canalizing functions* (NCFs) [15]. For example, a gene might be activated whenever one of multiple transcription factors is present - an example of a nested-canalizing OR function. This raises multiple important questions for the dynamical analysis of Boolean GRNs:

1. How much do the rules in expert-curated Boolean GRN models differ from threshold rules?

2. How different are the dynamics of expert-curated GRN and corresponding threshold networks?

3. There exist multiple threshold formalisms. Is one consistently “better” than another?

Using the repository of 122 expert-curated Boolean GRNs as the closest proxy of a “ground truth”, this manuscript attempts to answer these questions.

## 2 Methods

### 2.1 Boolean networks

A *Boolean network F* = (*f*_1_, …, *f*_*n*_) constitutes a popular modeling framework in systems biology. It consists of *n* nodes (e.g., genes, proteins, etc.). Each node can only be in two states, denoted 𝔽_2_. Commonly, 𝔽_2_ = {0, 1} or 𝔽_2_ = {−1, 1}. With biology in mind, we refer to the two states as absence or OFF (e.g., low protein concentration) and presence or ON (e.g., high protein concentration), respectively. Each node *x*_*i*_ in a Boolean network possesses a Boolean *update function* 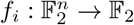, which describes the state of *x*_*i*_ at the next time point given the current state of the system. Under a synchronous updating scheme, all nodes are updated simultaneously. In this case, 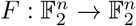 defines a deterministic state transition graph, also known as *state space*, which consists of 2^*n*^ states 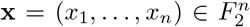 and their deterministically defined transitions. Asynchronous updating schemes allow for nodes to be updated separately, at potentially different time scales [16]. In most schemes, only a single node is updated at a time. In this case, the state space is typically stochastic because 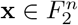 may transition to *n* different states 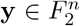, depending on which node *x*_*i*_ is updated. In this study, we consider both a synchronous updating scheme and a general asynchronous updating scheme (each node is updated with equal probability [17]), which has been established as the most efficient and informative asynchronous updating method [18].

### 2.2 Boolean threshold functions

A Boolean function is a threshold function if there exists a hyperplane that separates the points where the function is OFF and ON. More precisely, 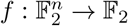 is a threshold function if there exist weights *w*_1_, …, *w*_*n*_ ∈ ℝ and a threshold *θ* ∈ ℝ such that for all 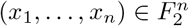,

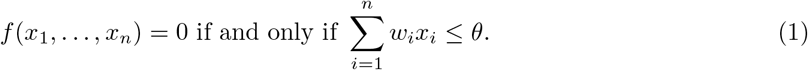

Threshold functions arise in a number of applications ranging from electrical engineering to reliability and game theory [19]. Threshold functions are also frequently used in GRN inference. Here, only the type of regulation (activation vs inhibition) is typically known from experiments. In other words, the sign of the weights *w*_*i*_ is known but the magnitude is unknown. Thus, all standard threshold formalisms used in GRN inference assume, in the absence of further experimental evidence, *w*_*i*_ ∈ {−1, 1}, *θ* = 0. We refer to such threshold functions as *basic*. The weights *w*_*i*_ can be thought of as labels of the edges of the wiring diagram of a biological Boolean network. That is, all experimental evidence is contained in the *wiring diagram W* ∈ {−1, 0, 1}^*n*×*n*^, also known as *interaction matrix*, where

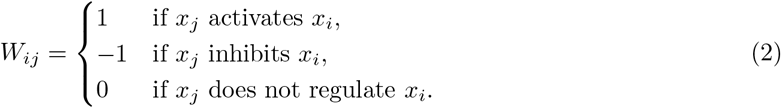

In this manuscript, we assume that the wiring diagram has already been inferred and compare two threshold formalisms that are frequently employed to model the dynamics of biological Boolean networks: the Ising and the 01 formalism. The Ising formalism stems from statistical physics where it describes the zero temperature Glauber dynamics of a disordered Ising model, commonly used to model ferromagnetic hysteresis in ferromagnets and spin glasses [20–22]. In the Ising formalism, the two binary states are 𝔽_2_ = {−1, 1}. In biological applications, −1 corresponds to the absence of a protein (i.e., OFF) and 1 to the presence (i.e., ON). In the 01 formalism, the two binary states are instead F_2_ = {0, 1}, with 0 indicating the OFF state. For both threshold formalisms, the next state of node *x*_*i*_ is derived from the current state of the network as follows:

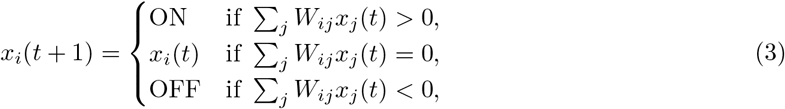

where *W* ∈ {−1, 0, 1}^*n*×*n*^ is the underlying wiring diagram.

For an alternative form of Equation 3, we define *A*_*i*_, *I*_*i*_ ⊆ {1, 2, …, *n*} as the set of indices of activators and inhibitors of node *x*_*i*_. That is, *A*_*i*_ = {*j* : *W*_*ij*_ = 1} and *I*_*i*_ = {*j* : *W*_*ij*_ = −1}. With this, we have

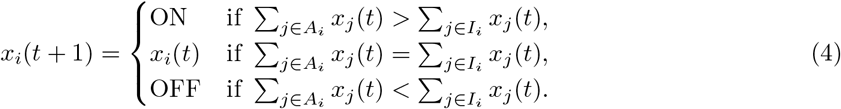

Whether the OFF state is represented by −1 or 0 has a major implication: A 01 threshold function is ON if the number of present (i.e., ON) activators is higher than the number of present inhibitors. If both numbers are equal, the state of node *x*_*i*_ does not change. The number of absent (i.e., OFF) activators and inhibitors does not matter in the 01 formalism but it does matter in the Ising formalism. An Ising threshold function is ON if the number of present activators and absent inhibitors is higher than the number of absent activators and present inhibitors.

For ease of comparison, we solely use 𝔽_2_ = {0, 1} throughout the Results and Discussion. To express both 01 *and* Ising threshold functions as Boolean functions *f* : {0, 1}^*n*^ → {0, 1}, we modify Equation 4 slightly by replacing OFF and ON with 0 and 1. We define *X*^0^(*t*) = {*j* : *x*_*j*_(*t*) = 0} and *X*^1^(*t*) = {*j* : *x*_*j*_(*t*) = 1}. With this, the 01 threshold formalism yields

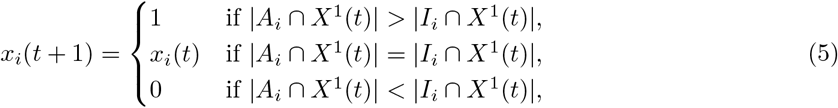

while the Ising formalism yields

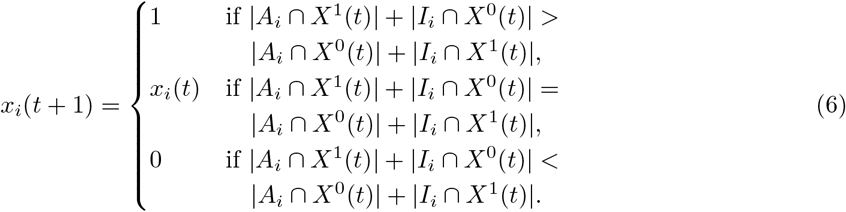

### 2.3 Similarity between Boolean functions

To assess the similarity between two Boolean functions *f* (*x*_1_, …, *x*_*m*_) and *g*(*x*_1_, …, *x*_*n*_) where *m*≤ *n*, we generate an extended Boolean function 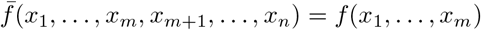, which does not depend on the last *n* − *m* variables. We then quantify the similarity *s*(*f, g*) between *f* and *g* as the proportion of inputs *x* = (*x*_1_, …, *x*_*n*_) ∈ 𝔽_2_ for which 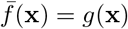. That is,

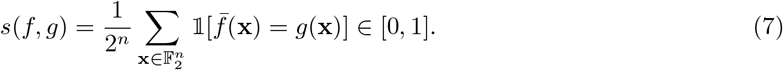

For example, to compare *f* (*x*_1_) = *x*_1_ and *g*(*x*_1_, *x*_2_) = *x*_1_ ∨ *x*_2_, we generate 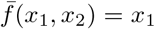 and observe that 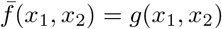 whenever (*x*_1_, *x*_2_) ≠(0, 1). Since *f* and *g* yield the same output for 3 out of 4 possible inputs, *s*(*f, g*) = 3*/*4.

### 2.4 Similarity between Boolean network dynamics

We employ several measures to quantify the similarity between the dynamics of two Boolean networks 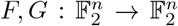. The state space describes the entire network dynamics. The overlap between the synchronous state spaces of networks *F, G*, denoted *𝒟 𝒮* _sync_(*F, G*), can be quantified by the proportion of states that update to the same state under *F* and *G*. I.e.,

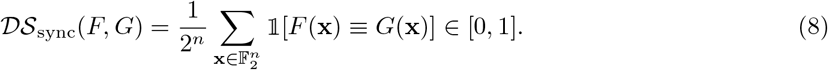

Instead of the indicator function, we can use the mean to quantify the difference between *F* (**x**) and *G*(**x**). This yields an alternative measure

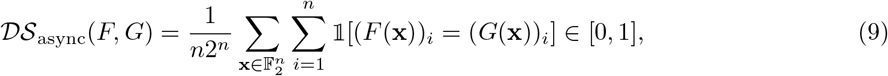

which describes the overlap between the state spaces of the two networks under the general asynchronous updating scheme. With Eq. 7, this is simply the mean similarity of the Boolean functions, i.e.,

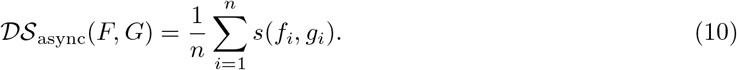

Due to their finite size and deterministic transitions, synchronously updated Boolean networks eventually exhibit periodic dynamics and settle into an attractor – either a steady state or a limit cycle. Asynchronously updated Boolean networks – with stochastic transitions – share the same steady states but they rarely have periodic limit cycles. Instead, they often possess so-called complex attractors, a set of states wherein the system oscillates, typically without any periodicity. In biological networks, the attractors correspond to phenotypes or cell types. To assess the similarity of the attractor spaces of Boolean networks *F, G*, we first obtain the network attractors and corresponding basin sizes. While the entire state space of small Boolean networks can be computed quickly, the exponentially-increasing size of the state space makes this impossible for large networks. To avoid introducing bias, we used the same method to sample the state space of each network, irrespective of network size. For each network, we generated *N* = 1, 000 random initial states and repeatedly updated each initial state until it transitioned into a periodic orbit, associated with an attractor. The proportion of random initial states that transitioned to a specific attractor served as estimate for the corresponding relative basin size. We note that an attractor with relative basin size *b* ∈ (0, 1] will be found with probability *p*(*b, N*) = 1 − (1 − *b*)^*N*^. For example, we will identify attractors with relative basin sizes of *b* = 1% and *b* = 0.1% with a probability of 99.996% and 63.23%, respectively. Thus, while our sampling approach cannot reliably identify all attractors, it will find all attractors with a decent basin size (i.e., all biologically meaningful attractors) and provides unbiased estimates of the relative basin sizes.

Let the lists *𝒜* (*F*), *𝒜* (*G*) describe the attractors of *F* and *G*. Each element in these lists is an ordered sequence of states (**x**^(1)^, …, **x**^(*l*)^) where *l* is the length (i.e., periodicity) of the attractor. Further, let *ℬ* (*F*) ∈ (0, 1]^| *𝒜* (*F*)|^, *ℬ* (*G*) ∈ (0, 1]^| *𝒜* (*G*)|^ describe the relative basin sizes corresponding to the attractors. Note that ∑*b*∈*B*(*F*) *b* = ∑*b*∈*B*(*F*) *b* = 1. As an example, consider the 2-node Boolean network *F* (*x*_1_, *x*_2_) = (*x*_1_ ∨ *x*_2_, *x*_1_ ∧ ¬*x*_2_). Under synchronous update, *F* has two attractors, a steady state and a 2-cycle: *𝒜* (*F*) = [(00), (10, 11)]. The state 01 transitions to 10, which is part of the 2-cycle. Thus, *ℬ* (*F*) = [1*/*4, 3*/*4]. From the set of attractors and corresponding basin sizes, we can compute the long-term distribution *µ*_*F*_ of a Boolean network *F* when initializing *F* at random (i.e., when starting from any state 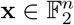 with equal probability). For any 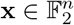, we have

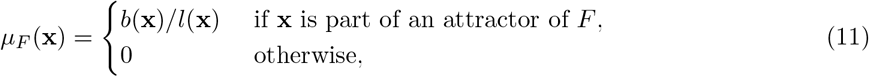

where *b*(**x**) is the basin size and *l*(**x**) is the length of the attractor that contains **x**. By design, ∑**x** *µ*_*F*_ (**x**) = 1. We can then compute the Jensen-Shannon distance metric [23] between *µ*_*F*_ and *µ*_*G*_ to assess the similarity of the attractor spaces of Boolean networks *F, G*,

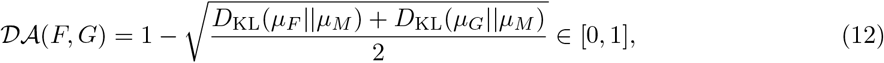

where *D*_KL_ is the Kullback-Leibler divergence and *M* = (*µ*_*F*_ + *µ*_*G*_)*/*2 is the mixture distribution (i.e., the pointwise mean) of *µ*_*F*_ and *µ*_*G*_.

Biological Boolean network models contain a surprisingly high number of so-called source nodes, which remain constant at all time and typically codify a cellular context or external inputs [15,24]. A network with *k* source nodes has at least 2^*k*^ attractors. To avoid any confounding effect of the number of source nodes on the network-level similarity metrics, we computed *𝒟 𝒮*1(*F, G*), *𝒟 𝒮*2(*F, G*) and *𝒟 𝒜* (*F, G*) only for “fixed-source” networks (as described in detail in [24]), and reported the average across all sampled fixed-source networks. In short, for a Boolean network with *k* source nodes, we randomly selected min(32, 2^*k*^) different source node states and replaced all instances of *n* in the formulas by the number of nodes that are not source nodes. We note that this approach only works as long as *F* and *G* have the same source nodes, which is the case in our study, in which *F* and *G* even possess the same exact wiring diagram.

### 2.5 Repository of expert-curated GRN models

The largest repository of expert-curated Boolean GRN models contains 122 distinct models [15]. Here, “distinct” means that the variables of any two of the 122 models overlap *<* 90%. Threshold rules can only be generated for a GRN with defined directionality and signs (activating vs inhibiting). Fifteen models contain regulators that have a conditional effect (i.e., both activating and inhibiting depending on the state of the other regulators). We excluded these regulatory functions (0.9% of all 5,112 regulatory logic functions) from the function-level analyses, reported in Figs. 4,5,6. For all network-level analyses, reported in Figs. 7,8,9, we analyzed 100 of the 122 distinct models. In addition to the 15 models with conditional regulators, we excluded seven models, which possess regulatory functions with 15 or more inputs (exclusion for computational reasons). For all presented analyses, we derived from each published model its signed wiring diagram W (Equation 2), which we never modified. In other words, we assumed the signed wiring diagram associated with a published model is 100% correct.

**Figure 4.**
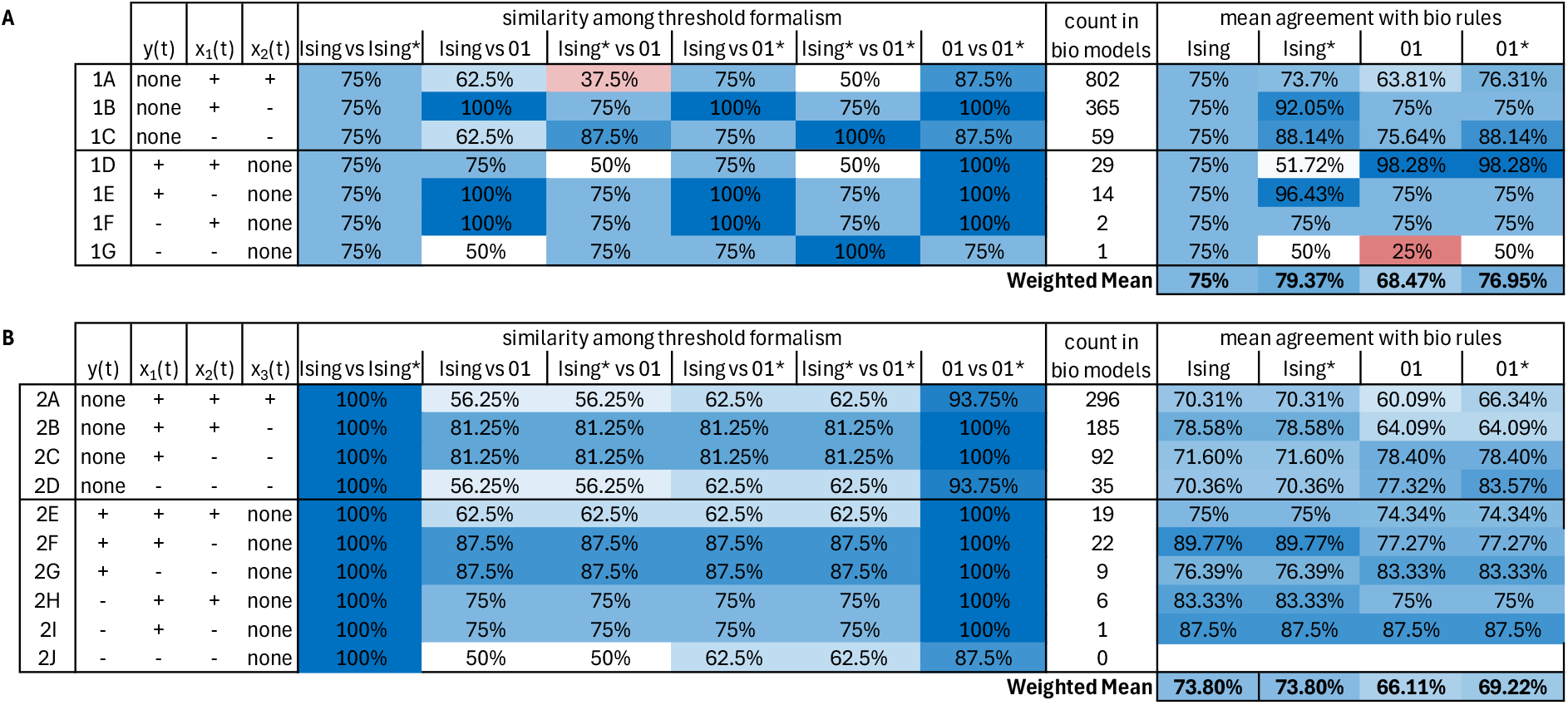
Similarity between different types of threshold rules and published biological rules for degree (A) 2 and (B) 3. The leftmost column contains a reference to Figures 2 and 2. The next columns describe the type of regulation of each input: positive (+) and negative (−). The expected similarity among two Boolean functions is always 50%. Higher-than-expected (lower-than-expected) values are colored in shades of blue (red).

**Figure 5.**
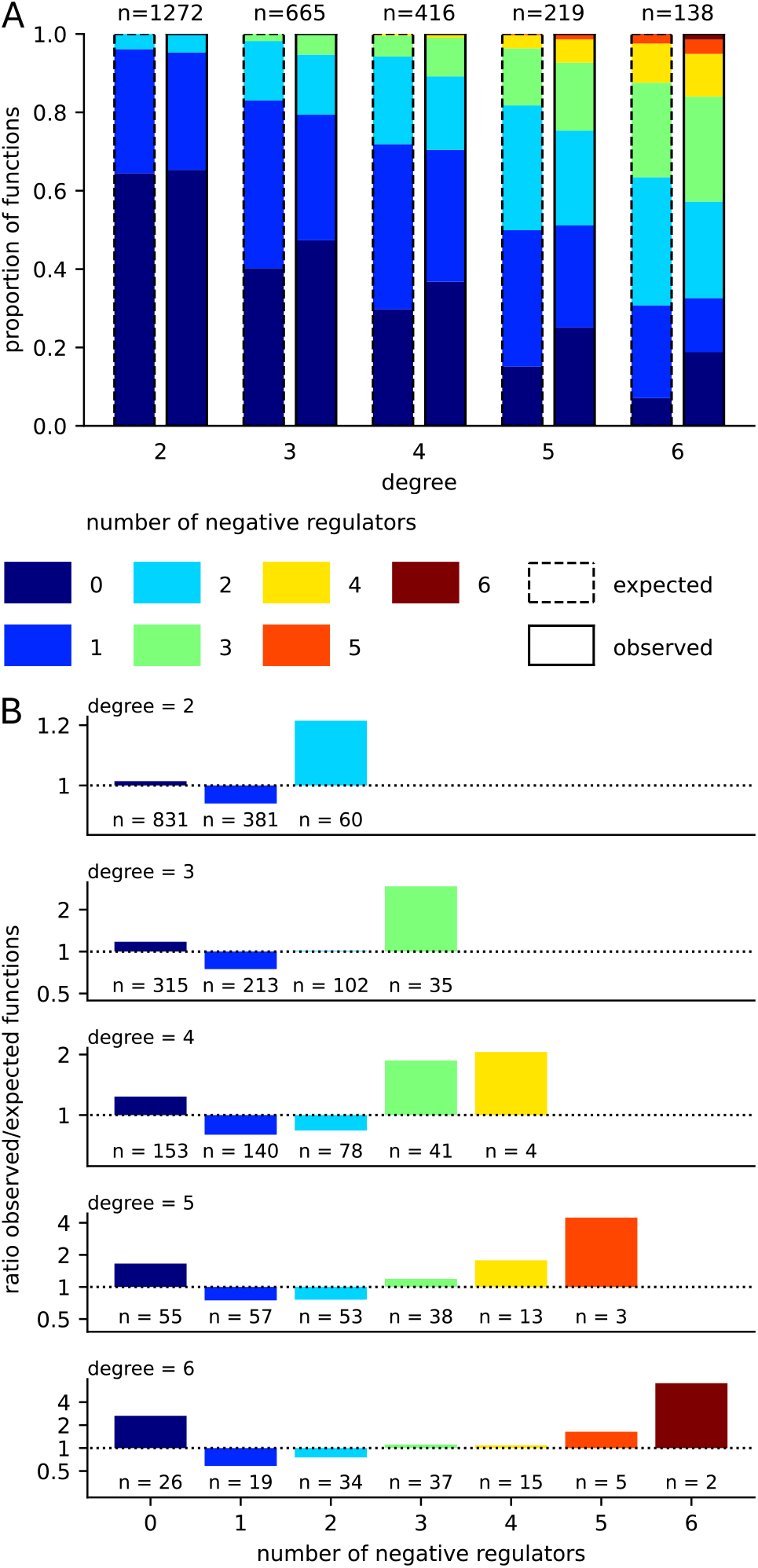
Boolean biological regulatory rules tend to have regulators of the same type. (A) Stratification of all expert-curated Boolean biological regulatory rules based on the number of inputs (x-axis) and the number of negative inputs (color). Rules that contain conditional inputs are excluded. Each observed distribution (the right bars with solid borders) is compared to the expected distribution (the left bars with dashed borders), which is computed based on the proportion of positive vs negative inputs for functions of a given degree and the combinatorial likelihood. n = observed total number of biological rules of a given degree and without any conditional input. (B) Ratio of the observed vs expected proportion of functions with a given number of total inputs (sub panels) and negative inputs (x-axis). Ratios above 1 indicate types of functions that are enriched in expert-curated Boolean biological network models. n = observed number of biological rules of a given number of total inputs and negative inputs, excluding rules with any conditional input.

**Figure 6.**
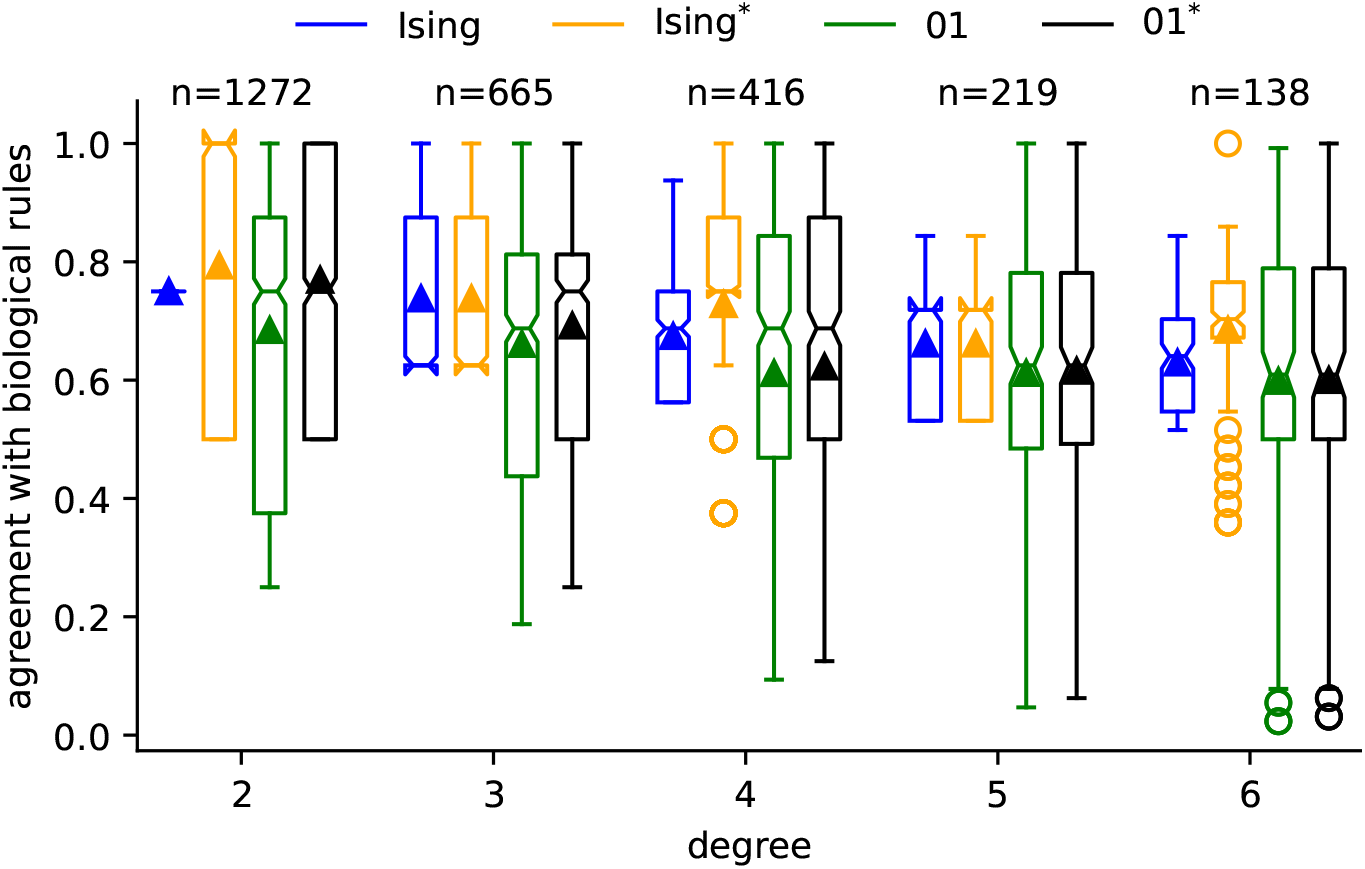
Similarity between threshold rules and published biological rules. All published biological rules of a given degree (x-axis) and without any conditional regulators are compared to different threshold rules (color). Each box extends across the interquartile range (IQR), the notch shows the median, the triangle shows the mean, whiskers extend from the box to the farthest data point lying within 1.5 IQR from the box, and circles (outside the whiskers) show outliers.

**Figure 7.**
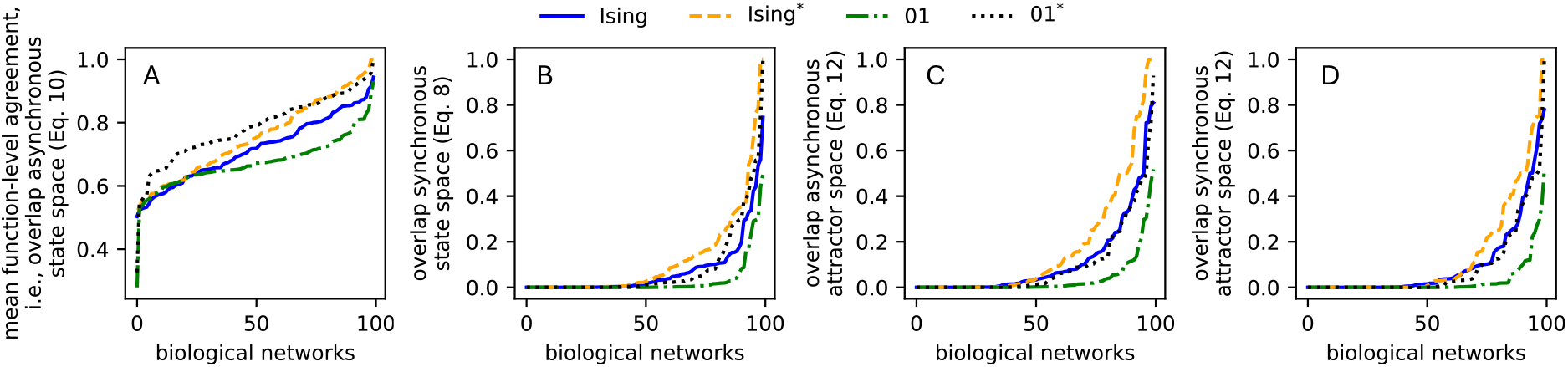
Ability of threshold networks to recover the dynamics of biological networks. For four threshold formalisms (color) and four dynamic similarity measures ((A) the mean agreement of the regulatory functions (i.e., the overlap of the asynchronous state space), see Eq. 10, (B) the overlap of the synchronous state space, see Eq. 8, and (C-D) the overlap of the (C) asynchronous and (D) synchronous attractor space, see Eq. 12), the empirical cumulative density function across the 100 biological networks is shown.

**Figure 8.**
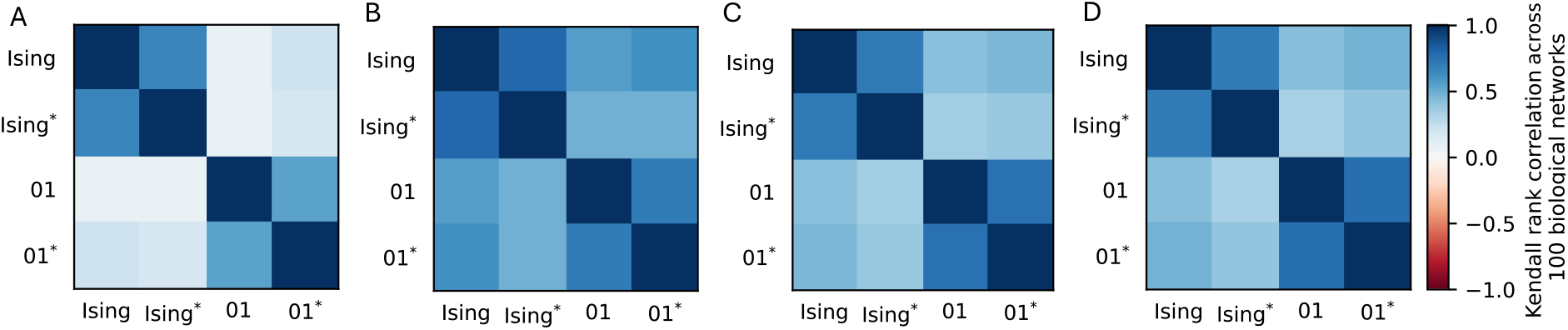
Pairwise agreement among threshold formalisms in dynamic similarity rankings of biological networks. For four threshold formalisms, the dynamic similarity ((A) the mean agreement of the regulatory functions (i.e., the overlap of the asynchronous state space), see Eq. 10, (B) the overlap of the synchronous state space, see Eq. 8, and (C-D) the overlap of the (C) asynchronous and (D) synchronous attractor space, see Eq. 12) with 100 biological networks is computed, see Fig. 7. The heatmaps display the similarity (quantified using Kendall’s *τ* coefficient) of the rankings of the biological networks between any two threshold formalisms.

**Figure 9.**
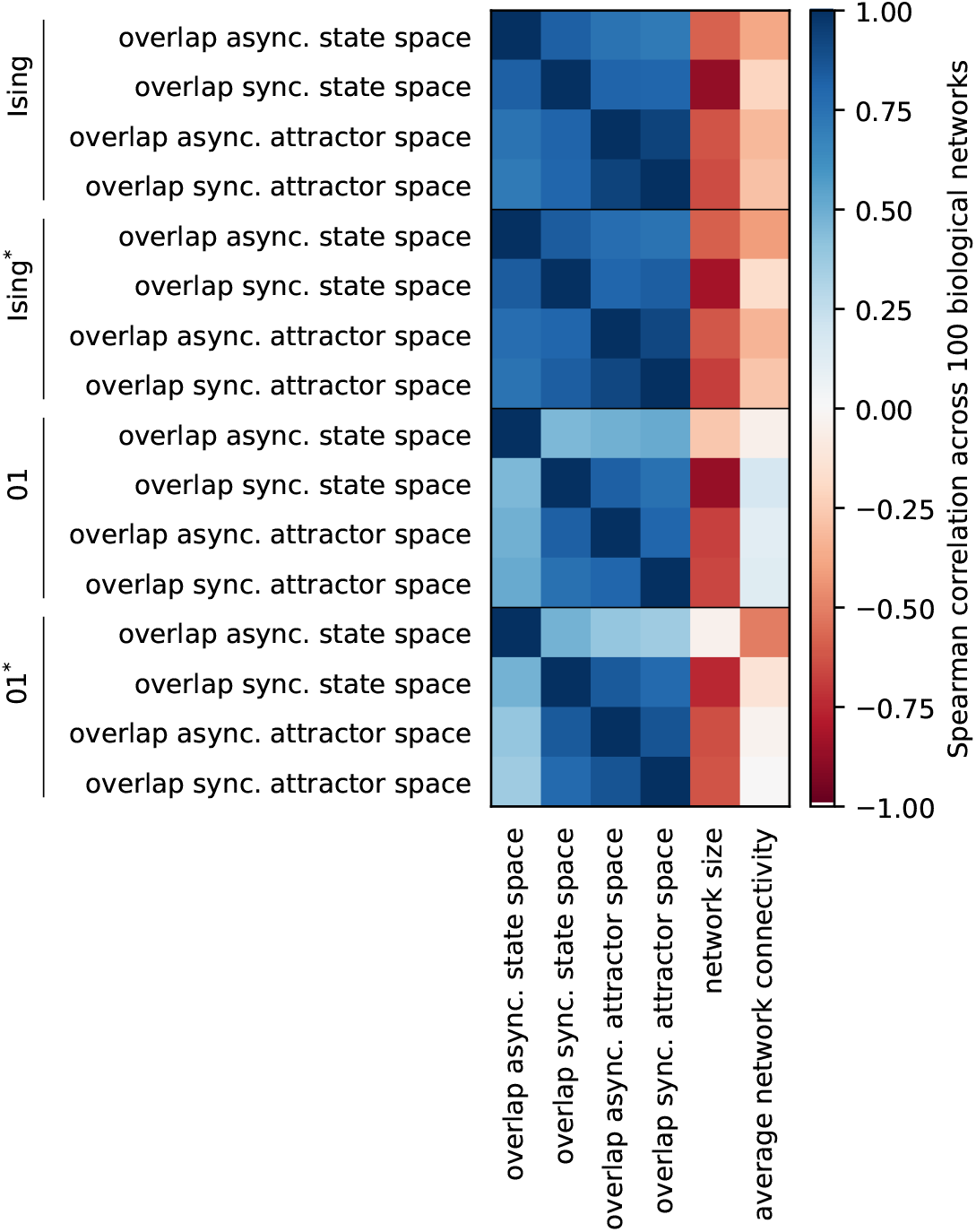
Correlation among dynamics similarity measures and network properties. For each threshold formalism (4 rows each), the Spearman correlation of dynamics similarity measures with other dynamics similarity measures, as well as network properties, is computed across the 100 biological networks.

## 3 Results

### 3.1 Similarity between threshold formalisms

Throughout, we compare two different threshold formalisms: the Ising formalism and the 01 formalism. We begin with a theoretical analysis and a comparison of the two types of threshold rules. Ising threshold rules are ON (OFF) if the number of present activators and absent inhibitors is higher (lower) than the number of absent activators and present inhibitors. Thus, Ising rules are, by design, always unbiased, i.e., both binary values, ON and OFF, occur with the same probability when considering the rule in truth table format. On the other hand, a 01(−threshold) rule is ON (OFF) if the number of present activators is higher (lower) than the number of present inhibitors. The number of absent regulators does not affect 01 rules. Thus, 01 rules are only unbiased when the number of activators and inhibitors is the same.

To compare the differences in threshold formalisms more comprehensively, we considered all possible Boolean functions *y*(*t*+1) = *f* (*x*_1_(*t*), *x*_2_(*t*)) with two inputs (Fig. 2A-C) and *y*(*t*+1) = *f* (*x*_1_(*t*), *x*_2_(*t*), *x*_3_(*t*)) with three inputs (Fig. 3A-D). Note that *y* may regulate itself, in which case we have *y*(*t*+1) = *f* (*y*(*t*), *x*_1_(*t*)) (Fig. 2D-G) and *y*(*t* + 1) = *f* (*y*(*t*), *x*_1_(*t*), *x*_2_(*t*)) (Fig. 3E-J). Since each input in a threshold rule represents either an activator or an inhibitor, there are seven (ten) possibly different basic threshold functions with two (three) inputs. If the number of activators equals the number of inhibitors, Equation 5 and Equation 6 are equal. This is the only case where the Ising formalism and the 01 formalism yield the same Boolean functions (Fig. 2B,E,F). Whenever there are strictly more (fewer) activators than inhibitors, 01 rules contain more (fewer) ones than zeros and are biased (Fig. 2A,C,D,G and Fig. 3). It is therefore not very surprising that Ising rules and 01 rules differ most if all regulators are of the same type (Fig. 4).

In the case of auto-regulation, all investigated threshold formalisms can give rise to degenerated functions, which do not depend on all its inputs. For example, the auto-regulatory 2-input threshold function *y*(*t* + 1) = *f* (*x*(*t*), *y*(*t*)) simplifies to *y*(*t*) when *y* is an activator and *x* is an inhibitor (Fig. 2E), while it simplifies to *x*(*t*) if *x* is an activator and *y* and inhibitor (Fig. 2F). This is the case for both threshold formalisms. Under the Ising formalism, *f* (*x*(*t*), *y*(*t*)) also simplifies to *y*(*t*) if *x* and *y* are both activators, completely disregarding the regulatory effect of *x* (Fig. 2D). The corresponding 01 rule is *f* (*x*(*t*), *y*(*t*)) = *x*(*t*) ∨ *y*(*t*). If both *x* and *y* are inhibitors, then *f* (*x*(*t*), *y*(*t*)) = ¬ *x*(*t*) under the Ising formalism and *f* (*x*(*t*), *y*(*t*)) = 0 under the 01 formalism (Fig. 2G). An analysis of auto-regulatory Boolean 01 threshold functions *y*(*t* + 1) = *f* (*x*_1_(*t*), …, *x*_*n*−1_, *y*(*t*)) with degree *n* ≥ 2 revealed that the value of *y*(*t*) does not at all affect *f* if and only if *y*(*t*) is an inhibitor (see e.g., Fig. 3E-J). This fact can be proven mathematically by expressing 01 threshold rules as

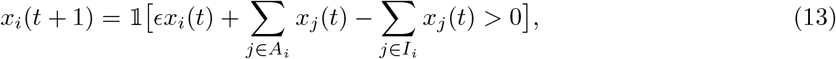

which is equivalent to Equation 4 for any *ϵ* ∈ (0, 1).

In the case of inhibitory auto-regulation (i.e., *i* ∈ *I*_*i*_),

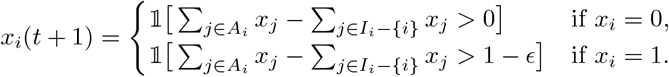

Since *x*_*j*_ ∈ {0, 1}, the two cases are always equal, which proves that a 01 threshold rule never depends on an inhibitory auto-regulatory input.

In the case of activating auto-regulation (i.e., *i* ∈ *A*_*i*_),

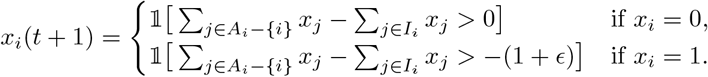

Here, the value of the activating auto-regulatory input *x*_*i*_(*t*) matters in determining *x*_*i*_(*t* + 1). The gap of 1 + *ϵ* between the two cases proves that *x*_*i*_(*t*) is even more important than the other inputs (i.e., *x*_*i*_ has higher activity [25] and higher edge effectiveness [26]), unless all inputs are activators. This can be seen in Figs. 2E, 3F,G.

Using the same arguments, we can show that the same is true for auto-regulatory Ising rules, however only if their degree is even. If the degree is odd, an Ising rule is always determined by the state of its activators and inhibitors because equality in the conditions of Equation 6 cannot occur. These observations likely explain why some expert-curated Boolean regulatory logic rules contain non-essential variables [15].

### 3.2 Insights from expert-curated regulatory logic functions can inform improved threshold formalisms

The largest repository of expert-curated biological Boolean network models consists of 122 distinct models with a total of 5112 regulated nodes [15]. Prior analysis of the 5112 Boolean update rules has revealed that biological regulatory functions are more canalizing, more redundant and more biased than expected [15]. Moreover, 73.9% of regulators have a positive (i.e., activating) effect on the target node, i.e., an increased expression of the regulator (that is, a change from 0 to 1) leads to an increased expression of the target for some states of the other regulators, and possibly no change in the target for other states of the other regulators. 23.6% of regulators have a negative (i.e., inhibiting) effect, and 0.9% of regulators have a conditional effect (i.e., both activating and inhibiting depending on the state of the other regulators). Further, the higher the number of regulators the lower is the proportion of activating regulators: Excluding functions with conditional regulators, the proportion of inputs that are activators is *p*_*a*_(*n*) = 78.6%, 71.3%, 71.0%, 65.1%, 61.8% in biological regulatory functions with degree *n* = 2, 3, 4, 5, 6, respectively. This implies that the probability that an n-input function contains *a* activators and *n* – *a* inhibitors is 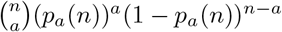. Interestingly, we found that biological regulatory functions tend to have regulators of the same type (Fig. 5). That is, more genes than expected are regulated only by activators, and many more genes than expected are solely inhibited. On the contrary, genes that have exactly one inhibitor proved particularly rare. These trends were consistent across all investigated degrees (i.e., all degrees with sample size ≥ 100).

Most expert-curated GRN models govern cellular processes in eukaryotes [15]. The observed higher proportion of activating regulators can be contributed to the fact that eukaryotic genes are by default off and are typically only transcribed when needed [27]. This also explains another observation: published regulatory functions, especially those with many inputs (i.e., higher complexity), contain more 0s than 1s (in truth table format). The mean bias across all expert-curated functions with degree 2, 3, …, 6 is 45.64%, 43.71%, 39.42%, 38.51%, 35.2%, respectively.

In light of these observations, we propose a slight modification to each threshold formalism that defines the function value in some/all cases of equality among positive and negative forces in Equation 4. In the modified Ising formalism (denoted Ising^*^), the Boolean function is 0 (rather than remaining at its current value) in the case of equality among positive and negative forces in Equation 6. That is, we assume the default state of a gene to be off, the standard for eukaryotic genes. The modified Ising formalism thus takes the following form:

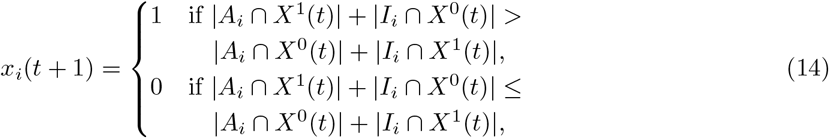

where *A*_*i*_, *I*_*i*_, *X*^0^(*t*) and *X*^1^(*t*) are defined as in Eq. 6.

If a gene is only regulated by inhibitors, the 01 formalism yields strange Boolean functions: the zero function in the presence of auto-regulation (Fig. 2G, Fig. 3J), and, in the absence of auto-regulation, a function that prevents a gene from ever becoming expressed again once it is not expressed (Fig. 2C, Fig. 3D). We therefore propose the following modification (denoted 01^*^): If a gene is only regulated by inhibitors and all inhibitors are absent, the 01^*^-function is 1 (i.e., the gene is expressed). Likewise, if a gene is only regulated by activators and all activators are absent, the 01^*^-function is 0 (to ensure such a gene does not remain expressed forever). The modified 01 formalism takes the following form:

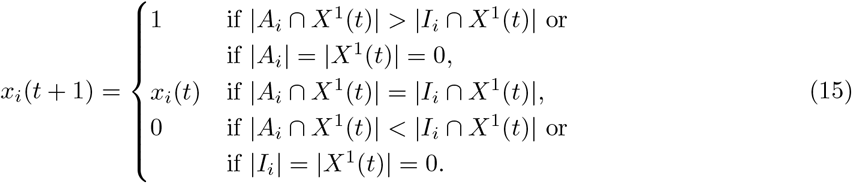

### 3.3 Differences between biological regulatory logic and threshold rules

Next, we compared the similarity between threshold rules and biological rules. All 2-input biological rules *f* (*x*_1_, *x*_2_) are nested canalizing [15,28]. That is, they are of the form *f* = (¬)*x*_1_ ∨ (¬)*x*_2_ or *f* = (¬)*x*_1_ ∧ (¬)*x*_2_ where a negation appears whenever *x*_*i*_ is an inhibitor. For any such function, the bias (i.e., the proportion of 1s in truth table format) is 1*/*4 = 25% or 3*/*4 = 75%. This explains why 2-input Ising rules (with bias 50%) agree with biological rules for 75% of the input combinations, irrespective of the type of the two inputs (Fig. 4A). On the other hand, the average agreement between 01 threshold and biological rules depends on the type of the inputs. If *x*_1_ and *x*_2_ are both activators, with *n* = 802 the most frequently observed case, 01 rules only agree with biological rules on average for 63.81% of input combinations. If *f* = *x*_1_ ∨ *x*_2_, the agreement is high with 7*/*8 = 87.5%. However, if *f* = *x*_1_ ∧ *x*_2_, the agreement is only 3*/*8 = 37.5%. By forcing *f* (*x*_1_ = 0, *x*_2_ = 0) = 0 in the modified 01 rule, the agreement increases by exactly 1*/*8 = 12.5% to an average of 76.31%. Overall, the modified Ising rules exhibit the highest average agreement with 2-input biological rules (79.37%), followed by modified 01 rules (76.95%), Ising rules (75%) and 01 rules (68.47%). For degree *n* = 3, Ising rules (and modified Ising rules which do not differ from Ising rules whenever the degree is odd) match biological rules better than modified 01 rules and 01 rules with a mean agreement of 73.80%, 69.22%, 66.11%, respectively (Fig. 4B). The same is true for degree *n* ∈ {4, 5, 6} (Fig. 6). The higher the degree the smaller is the average agreement between threshold and biological rules, irrespective of the choice of threshold formalism. Moreover at higher degrees, the differences between 01 and modified 01 rules become negligible (because the maximal possible difference between the two rules is 1*/*2^*n*^, which decreases as the degree *n* increases), while differences among Ising and modified Ising rules, as well as differences in their agreement with biological rules persist. As already described above for the case of two activators, 01 rules match some nested canalizing biological rules almost perfectly – modified 01 rules even equal some biological rules (Table 1). However, the agreement with other biological rules is rather low.

**Table 1.** Bias and similarity with corresponding threshold functions for the most frequent 3-input NCFs in published GRN models. NCFs with the same layer structure [29] and the same number of activators per layer (second column) are equivalent as they have the same bias and similarity with corresponding threshold functions.

| Function f | Layers and types | count in bio models | bias |  |  |  | Similarity of f and |  |  |
| --- | --- | --- | --- | --- | --- | --- | --- | --- | --- |
|  |  |  | f | Ising(f) | 01(f) | 01*(f) | Ising(f) | 01(f) | 01*(f) |
| $X \vee Y \vee Z$ | +++ | 117 | 7/8 | 1/2 | 15/16 | 7/8 | 5/8 | 15/16 | 1 |
| $X \wedge Y \wedge Z$ | +++ | 87 | 1/8 | 1/2 | 15/16 | 7/8 | 5/8 | 3/16 | 1/4 |
| $\neg X \ \& \ (Y \vee Z)$ | -, ++ | 73 | 3/8 | 1/2 | 11/16 | 11/16 | 7/8 | 11/16 | 11/16 |
| $X \ \& \ Y \ \& \ \neg Z$ | ++- | 58 | 1/8 | 1/2 | 11/16 | 11/16 | 5/8 | 7/16 | 7/16 |
| $X \ \& \ \neg Y \ \& \ \neg Z$ | +- - | 56 | 1/8 | 1/2 | 5/16 | 5/16 | 5/8 | 13/16 | 13/16 |
| $X \vee (Y \wedge Z)$ | +, ++ | 46 | 5/8 | 1/2 | 15/16 | 7/8 | 7/8 | 11/16 | 3/4 |
| $X \wedge (Y \vee Z)$ | +, ++ | 45 | 3/8 | 1/2 | 15/16 | 7/8 | 7/8 | 7/16 | 1/2 |
| $\neg X \ \& \ \neg Y \ \& \ \neg Z$ | --- | 23 | 1/8 | 1/2 | 1/16 | 1/8 | 5/8 | 15/16 | 1 |
| $\neg X \vee (Y \wedge Z)$ | -, ++ | 18 | 5/8 | 1/2 | 11/16 | 11/16 | 7/8 | 13/16 | 13/16 |

Theoretical considerations reveal how well any monotonic Boolean function *f* can match its corresponding threshold rule *t*(*f*): The key idea is to compare the bias of the two functions. Let *p*_*f*_, *p*_*t*(*f*)_ ∈ [0, 1] describe the bias of *f* and its corresponding threshold rule, respectively. Then, any difference in bias implies that the similarity between the two functions must satisfy

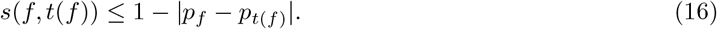

Specifically, *s*(*f, t*(*f*)) = 1 −| *p*_*f*_ − *p*_*t*(*f*)_ | if and only if the two functions match as closely as possible, i.e., if *f* = 0 whenever *t*(*f*) = 0 and *t*(*f*) = 1 whenever *f* = 1, when assuming, without loss of generality, that *pf* ≤ *pt*(*f*).

Ising rules are always unbiased, i.e., *p*_Ising(*f*)_ = 1*/*2, irrespective of *f*. The bias of 01-rules depends on the number of activating vs inhibitory inputs: As can be seen in Fig. 2A-C and Fig. 3A-D, a 01-rule with *n* non-auto-regulatory inputs, of which *m* ∈ {0, 1, …, *n*} are activating, has bias

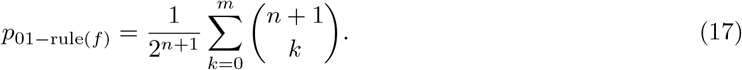

Similar formulas for the case of auto-regulation and modified 01-rules can be derived. These considerations explain why an AND-function *x*_1_ ∧ · · · ∧ *x*_*n*_ with low bias 1*/*2^*n*^ poorly matches its corresponding 01-threshold rule, which has bias 1− 1*/*2^*n*+1^, or why a NOT-AND-function ¬ *x*_1_ ∨· · ·∨¬ *x*_*n*_ with high bias 1 − 1*/*2^*n*+1^ poorly matches its 01-rule, which has bias 1*/*2^*n*+1^. Moreover, the results show that 01-rules are well-suited to describe the action of multiple independent transcription factors (which is modeled by an OR rule). However, if multiple activators form a protein complex, which regulates transcription, the appropriate biological rule is an AND function, which differs substantially from the 01-rule. This also explains why the deviations in agreement with biological rules are much larger for 01 rules than Ising rules, especially at higher degree (Fig. 6, Table 1).

### 3.4 Differences between the dynamics of biological networks and threshold networks

Thus far, we have only compared the similarity between expert-curated regulatory logic and corresponding threshold functions. The primary purpose of Boolean network models is however insight into the dynamic behavior of systems. Specifically, researchers are interested in the long-term behavior of systems as they settle into attractors, which in meaningful biological networks correspond to different cell types or phenotypes. We therefore asked the question: how well do threshold models recover (a) the entire synchronous state space (i.e., the entire dynamics) of a biological GRN, and (b) the attractors and corresponding basin sizes of a biological GRN?

To assess the similarity of two synchronous state spaces, we ask: Given a random state that is updated once according to each model, how similar are the updated states on average? If similarity of two states is quantified by the normalized Hamming distance (e.g., for **x** = (0, 0, 1, 1) and **y** = (0, 0, 0, 0), *d*(**x, y**) = 1*/*2), then the similarity of two synchronous state spaces is simply the average agreement of the regulatory functions, which we analyzed in Fig. 6. Using this measure of state space similarity, we found that modified 01 networks recovered the transitions in biological networks best (Fig. 7A), with an average similarity across the 100 biological networks of 78.36%, followed by modified Ising networks (75.44%), Ising networks (71.58%) and 01 networks (67.32%). Interestingly, the modified 01 formalism performed particularly well for networks with hard-to-recover state spaces (i.e., for networks with lower mean function-level agreement across the threshold formalisms). For networks that were easy to recover, the modified Ising formalism was slightly better. The similarity of two states can also be quantified by a binary indicator function (i.e., *d*(**x, y**) = 1 if **x** = **y** and 0 otherwise). As expected, this more stringent definition yielded much lower state space similarities (Fig. 7B). Across the 100 biological networks, modified Ising networks exhibited the highest state space overlap (mean = 12.46%), followed by modified 01 networks (8.56%), Ising networks (7.38%) and 01 networks (2.98%).

While a truthful representation of the entire state space is desirable, correct long-term behavior is arguably even more important. Boolean network trajectories eventually transition towards a network attractor. In biological networks, each attractor typically corresponds to a cell type or phenotype. Across the 100 biological networks, modified Ising networks had the highest average attractor space similarity (17.68% when asynchronously updated and 13.84% when synchronously updated), followed by Ising networks (11.41% and 10.14%), modified 01 networks (10.14% and 8.91%) and 01 networks (3.79% and 2.96%; Fig. 7C,D).

Fig. 7 displays an empirical cumulative density function across the 100 biological networks: For each threshold formalism, the networks are ranked in increasing order of dynamic similarity with corresponding threshold networks. This may imply that the dynamics of some biological networks are easier to recover than others, irrespective of the type of threshold formalism, which is not our intention. To quantify the degree to which the ranking in dynamic similarity differed among the threshold formalisms, we computed the pairwise Kendall rank correlation coefficient (Fig. 8). As expected, similar threshold formalisms (Ising and its modification, as well as 01 and its modification) mostly recover the same biological network dynamics well. On the other hand, biological networks with a higher function-level agreement with Ising and modified Ising rules did not exhibit any higher agreement with 01 rules and only slightly higher agreement with modified 01 rules (Fig. 8A). For the network-level dynamic similarity measures, all four threshold formalisms agreed to a substantial part on which biological network dynamics were easier to recover (Fig. 8B-D).

Thus far, we have ignored obvious confounders that affect how truthfully biological networks and their dynamics can be represented by threshold networks. These include the network size and the network connectivity. A Boolean network in *n* nodes has 2^*n*^ states. The larger the network the smaller is thus the chance that (a) a state is updated to exactly the same state, and (b) that attractors and their basins coincide in both the biological and the corresponding threshold network. This explains the observed strong negative Spearman correlations between network size and the overlap in state space and attractor space(Fig. 9). This trend was consistent across threshold formalisms. As shown before and summarized in Fig. 6, the average function-level agreement among biological and corresponding threshold rules decreases as the degree increases, irrespective of the threshold formalism. In line with this, more connected biological networks exhibited a lower average mean function-level agreement with their corresponding Ising, modified Ising and modified 01 threshold networks. Surprisingly, the mean function-level agreement of 01 networks with biological networks did not depend on the average network connectivity (*ρ*_Spearman_ = −0.04, *p* = 0.69). For Ising, modified Ising and 01 networks, network size was also significantly correlated with the mean function-level agreement (*ρ*_Spearman_ = − 0.59, − 0.6 and − 0.29, respectively). This is somewhat surprising, especially because network size and average connectivity are not significantly correlated across the 100 biological networks (*ρ*_Spearman_ = − 0.14, *p* = 0.16). Average network connectivity was weakly negatively correlated with the overlap of state space and attractor space for Ising and modified Ising networks but was not statistically significantly correlated for 01 and modified 01 networks. A weak negative correlation can be expected due to the reported decreasing agreement between biological and threshold rules as complexity increases.

## 4 Discussion

Boolean network models have long served as a cornerstone for investigating the dynamics of biological networks and specifically GRNs. In the absence of sufficient data to infer the exact regulatory logic, the Ising and 01 threshold formalisms are frequently used as default Boolean update rules. In our study, we critically evaluate these default models against a comprehensive repository of expert-curated Boolean GRN models, the closest available proxy of “reality”. By comparing their performance in recovering both the biological regulatory functions and the dynamic behaviors of cellular networks, our work addresses a key challenge in systems biology: the accurate inference of GRN dynamics from high-throughput data.

We described how the two formalisms can be expressed using the same equation (Equation 3), with the sole difference that the OFF state is represented by −1 in Ising rules and by 0 in 01 rules. When auto-regulation is present, both types of threshold rules can contain non-essential inputs, which means they fail to accurately represent available biological knowledge in a Boolean context. Informed by a meta-analysis of 122 expert-curated Boolean GRN models [15], we introduced a modified version of each threshold formalism, without loss of generality that is characteristic of threshold rules. The modified threshold rules consistently outperform the standard models. Notably, these improvements are reflected in a higher agreement with biological regulatory rules and a more faithful recovery of state space transitions and attractor landscapes – features that are essential for representing cellular phenotypes. Overall, the Ising formalism outperformed the 01 formalism, and modified Ising rules emerged as best default rules of Boolean GRN models from our systematic assessment. Given the generality of the modifications, further changes to threshold rules specific to individual contexts are expected to improve the ability of threshold rules to accurately capture the underlying dynamics. These findings underscore the importance of refining Boolean threshold models to better mirror the underlying biology, thereby enhancing their utility in the computational analysis of gene regulatory networks.

This study has some limitations. First, we treat the repository of 122 expert-curated Boolean GRN models as ground truth. Unfortunately, we cannot rule out that the diverse set of researchers that designed the published models may have had some shared preconceived notions and biases about what reasonable regulatory logic should look like. However, it is worth noting – and should partially help mitigate the potential impact of these biases on this study – that any Boolean models whose regulatory rules were assumed to follow some default formalism (e.g., threshold rules), rather than being informed by expert knowledge and biological experiments, were excluded from the repository from the beginning. Second, the modification of the Ising formalism was informed by the observation that most genes in expert-curated Boolean GRN models are more frequently off, the default state in eukaryotes. Using for validation the same set of rules that was used to inform this modification gives rise to a somewhat circular argument. While the improved function-level similarity of modified Ising rules is thus expected, it is assuring to see their improved ability to recover Boolean GRN dynamics and especially their phenotypes (i.e., attractors). That being said, the low number of expert-curated prokaryotic Boolean GRN models means it is impossible to properly evaluate the suitability of the modified threshold functions as default rules for prokaroytic regulatory logic.

Lastly, we consider the problem of Boolean GRN model inference as a two-step procedure: (i) infer the signed, directed wiring diagram from high-throughput data, (ii) infer the Boolean update rules that best match the data and agree with the inferred wiring diagram. In this manuscript, we only focus on the second step, i.e., we assume the inferred signed, directed wiring diagram represents the truth. In reality, the inferred wiring diagram itself may contain edges with higher and lower confidence or even errors. Access to information on the confidence in an edge and its sign could potentially result in more accurate default Boolean update rules. Moreover, considering Boolean GRN model inference as a one-step procedure, i.e., inferring the best Boolean network directly from high-throughput data, may harbor additional benefits, which are beyond the scope of this study.

Future work could explore additional modeling frameworks to identify robust default approaches for simulating GRNs. Extending Boolean models with multilevel logical rules—allowing for discrete states such as low, medium, and high for key regulatory nodes—could enhance the representation of graded expression patterns [30]. Incorporating memory-dependent dynamics, where node states are influenced by their own past states, may better capture temporal dependencies in gene regulation [31, 32]. Furthermore, investigating how structural properties of GRNs—such as degree distributions, fractal organization, or modularity—influence system dynamics could inform the selection of appropriate modeling frameworks [32]. Together, these approaches would improve the capacity to simulate GRN dynamics, especially under conditions of incomplete knowledge about the underlying regulatory rules.

## Author contributions

C.K.: Conceptualization, Methodology, Formal Analysis, Investigation, Software, Visualization, Writing - original draft, Writing - review and editing; K.H.: Conceptualization, Methodology, Writing - review and editing.

## Author declaration

The authors declare no competing interests

